# Human CODIS STR Loci Profiling from HTS Data

**DOI:** 10.1101/047225

**Authors:** Darrell O. Ricke, Martha Petrovick, Johanna Bobrow, Tara Boettcher, Christina Zook, James Harper, Edward Wack, Eric Schwoebel

## Abstract

Human DNA identification is currently performed by amplifying a small, defined set of short tandem repeat (STR) loci (e.g. CODIS) and analyzing the size of the alleles present at those loci by capillary electrophoresis. High-throughput DNA sequencing (HTS) could enable the simultaneous analysis of many additional STR and single nucleotide polymorphism (SNP) loci, improving accuracy and discrimination. However, it is necessary to demonstrate that HTS can generate accurate data on the CODIS loci to enable backwards compatibility with the FBI NDIS database. Sequencing can also detect novel polymorphisms within alleles that migrate with identical sizes by capillary electrophoresis, improving allele discrimination, and enhancing human identification analysis. All CODIS alleles from an individual can be amplified in a single, multiplex PCR reaction, and combined with additional barcoded samples prior to sequencing. A computational tool for allele identification from multiplexed sequence data has been developed. With longer-read-length platforms, 99.6% allele calling accuracy can be achieved. In the course of STR sequencing protocol development, 12 novel allele sequences have been identified for multiple loci. Sequencing STR loci combined with SNPs will enable new forensic applications.

## I. INTRODUCTION

Sequence differences in the human genome are a cornerstone in human identification and forensic applications. Current forensic standards have been developed using unlinked short tandem repeat (STR) polymorphisms. The FBI has defined 20 (previously 13) core STR loci for the Combined DNA Index System (CODIS) database as the current forensic standard in the United States. Current forensic protocols identify STR alleles by length of polymerase chain reaction (PCR) amplicons sized on capillary electrophoresis (CE) instruments. While highly effective and well accepted by the forensics community, these methods tend to be time consuming, expensive, and, in many cases, require manual evaluation of the results. Recent improvements in sequencing technologies, driven by broad market forces, might be coopted by the forensics community to significantly increase capabilities in forensic DNA analysis, and simultaneously decrease the cost of producing and analyzing that data.

The potential in this has not gone unnoticed by several groups. Pyrosequencing has been done on single STR loci using custom base dispensation order[2, 3]. Divne *et al.* [3] demonstrated the feasibility of using pyrosequencing for identification of 9 CODIS loci (plus Penta E). Fordyce *et al.* [1] sequenced 5 of the CODIS loci using the 454 platform. Bornman et al. [5] demonstrated sequencing of all of the CODIS loci using the Illumina GAIIx short-read technology, and treated the amplified samples the same as genomic DNA, fragmenting large amplicons containing the STR regions. With improvements in read lengths, Illumina has released the ForenSeq DNA signature prep kit.

Characterization of STR loci in whole genome sequencing results has been demonstrated [6], where several alleles were missed because of the lower coverage of STR regions. In addition, an extra allele was identified at D5S818 which was not present in the corresponding CE profile, most likely due to an error in assembling short reads that do not span the STR allele. This demonstrates that assembly of the correct STR allele sequences is dependent upon sequence read lengths. As sequencing technologies achieve longer sequence reads, it may be advantageous to shift to highly multiplexed panels with many additional loci while maintaining backwards compatibility with CODIS. All of these investigations support the idea that current generation HTS technologies have the potential to provide accurate STR profiles.

This study is designed to examine methods to improve amplicon production and multiplex balance, demonstrate the feasibility of automated allele calling software, and identify new permutations of STR alleles. The reliable sequencing of STR loci and accurate allele calling enables simultaneous backwards compatibility with existing DNA profile databases while expanding into new forensics capabilities such as inferring biogeographic ancestry, extended kinship and externally visible traits.

## II. MATERIALS AND METHODS

### A. Sample collection and isolation

Human DNA was obtained from buccal swabs under a protocol approved by the MIT Committee On the Use of Humans as Experimental Subjects (COUHES) review board. Cotton swabs were rubbed on the inside of both cheeks, and DNA was extracted using the Qiagen QIAamp DNA Investigator Kit (Qiagen Cat. no. 56504) protocol.

### B. PGM sequencing with AmpliSeq panel 87957

Buccal swabs were sampled from individuals using Bode swabs. Genomic DNA was isolated using the Qiagen Investigator Kit, eluted in 50uL of lowTE (0.1mM EDTA), then quantitated by Quant-iT dsDNA High-Sensitivity Assay kit from Invitrogen. The primer panel was designed for 182 amplicons in two pools. They were amplified using Ion Torrent AmpliSeq kit, according to manufacture’s protocol, including the secondary amplification and final libraries were eluted in 25uL. Sets were compared using an extended FuPa digestion (20min for each step), which may increase reads through homopolymer regions. Samples were then quantitated by Quant-iT (same as above), and subsequently diluted to the 20pmol (or ~1.2x10e7 molecules/uL) quantity suggested by Ion Torrent, using an average length of 275 base pairs for the calculation. Molecules/uL = (sample concentration[ng/uL] × 6.022 × 1023) / (656.6 × 109 × amplicon length [base pairs]). Amplicon libraries were then used in the 400bp HiQ Ion OneTouch Template PGM reaction kit, followed by the 400bp Ion Torrent HiQ PGM Sequencing kit.

### C. Roche 454 sequencing

Buccal swabs were sampled from individuals using either Bode or Epicenter swabs. Genomic DNA was isolated using the Qiagen Investigator Kit, eluted in 50uL of water, quantitated by Quant-iT dsDNA High-Sensitivity Assay (Invitrogen), and amplified using AmpliTaq Gold 360 (Applied Biosystems). Each 25uL reaction contained: 0.8uL Taq 360 Gold polymerase (4 Units), 2.5mM final MgCl2, 200uM final each dNTP (800uM total), and 400nM for each primer. Cycling conditions: 95C for 10min, then 95C for 30sec; 60C for 1min; 72C for 1min 27 more times; followed by 10 min 72C; 4C. Primer sequences were identical with PowerPlex-16 primers except for those loci shown in Table 1. PCR products were purified by Agencourt Ampure XP beads. Alterations to standard protocol were: A 1.28x reaction volume of beads was used; beads air-dried for 10min after EtOH wash, and samples were eluted off beads in 25uL of low TE (0.2mM EDTA, 10mM Tris-HCl, pH8). Samples were then quantitated, end repaired and dA-tailed using NEBNext Quick DNA Library Prep Master Mix Set for 454 kit from New England Biolabs, and Roche 454 Rapid Library adapters added using NEBNext Adaptor Ligation. Ligated products were Ampure purified, quantitated, then pooled in a 1 molecule/bead ratio, (ave. length = 271 base pairs). The rest of the sequencing was carried out as specified in the Roche 454 Sequencing Method Manual, except that 1/16 of the recommended amount of Amp primer was used.

**Table 1.**
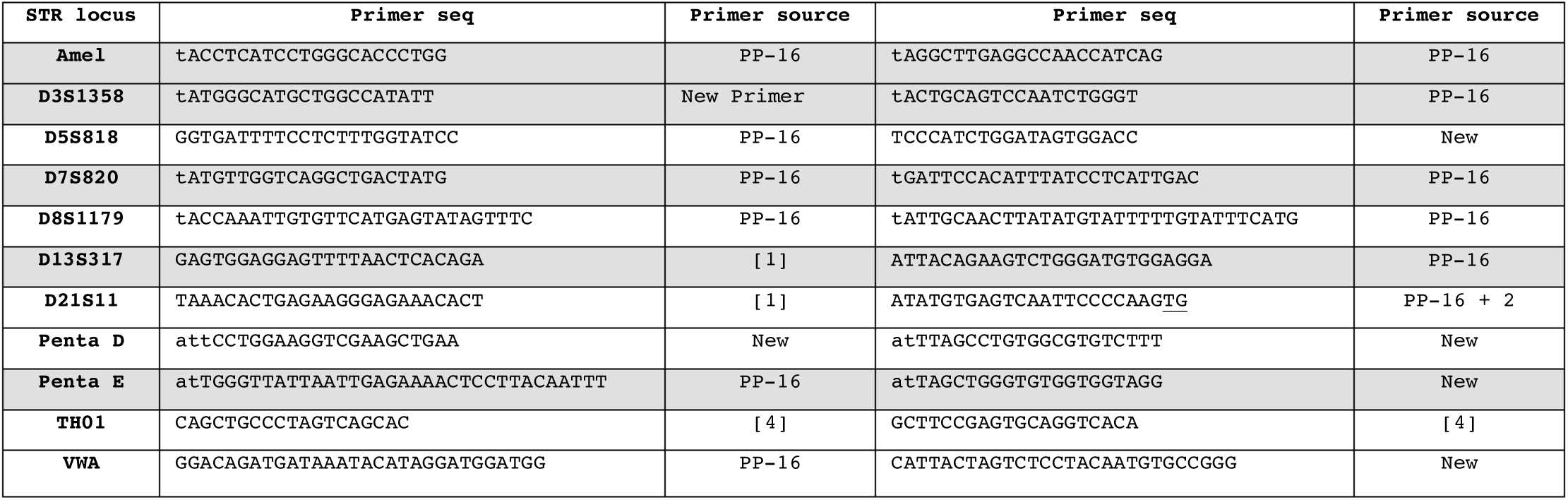
Primers used for balance-optimized multiplex amplification of STR loci. Upper case sequence indicates starting sequence. unmodified primers. Lower case sequence indicates additions to the 5’ end to improve NGS data balance. Underlined sequence indicates bases added to equalize Tm.

**Table 2.** Verified Novel STR Alleles Detected by Sequencing

| Locus | Allele | Allele sequence |
| --- | --- | --- |
| D3S1358 | 15" | TCTA[TCTG]<sub>1</sub>[TCTA]<sub>13</sub> |
| D5S818 | 10(11)(D4G->T) | [AGAT]<sub>11</sub> GT |
| D5S818 | 12(13) | [AGAT]<sub>13</sub> GT |
| D5S818 | 13(14)(U4T->C) | [AGAT]<sub>14</sub> GT |
| D8S1179 | 13' | [TCTA]<sub>13</sub> |
| D8S1179 | 14' | [TCTA]<sub>2</sub>[TCTG]<sub>1</sub>[TCTA]<sub>11</sub> |
| D8S1179 | 15' | [TCTA]<sub>2</sub>[TCTG]<sub>1</sub>[TCTA]<sub>12</sub> |
| D13S317 | 11' or 11(D1A->T) | [TATC]<sub>12</sub>AATC |
| D13S317 | 12' or 12(D1A->T) | [TATC]<sub>13</sub>AATC |
| D13S317 | 13' or 13(D1A->T) | [TATC]<sub>14</sub>AATC |
| D13S317 | 14' or 14(D1A->T) | [TATC]<sub>15</sub>AATC |
| vWA | 17'(19) | TCTA[TCTG]<sub>3</sub>[TCTA]<sub>13</sub> |

### D. Sanger sequencing

Sanger sequencing confirmed novel alleles that were identified by HTS sequencing. Amplicons were ligated into the TOPO pCR2.1 vector (Invitrogen catalogue number K203001) according to the manufacturer’s directions. Individual clones were purified from bacteria using the QIAprep Spin Miniprep kit (Qiagen catalogue number 27106) and submitted to Tufts University Core Facility for fluorescent Sanger sequencing.

### E. CE STR profiling

The amount of human DNA in the extracted DNA sample was quantified in the Quantifiler Duo DNA Quantification Kit (Applied Biosystems Cat. no. 4387746). Amplification using. Promega’s PowerPlex 16 kit was performed using 250-500 pg of template in a Veriti thermocycler, according to the manufacturer’s instructions. Separation and detection was carried out using an ABI 3130 with the injection parameters specified by Promega for the PowerPlex 16 profile, and the results were analyzed using Genemapper 3.2.

### F. Bioinformatics

The Ruby find_strs.rb program was developed to characterize multiplexed STR profiles. This program takes as input a FASTA file of DNA sequences, a barcode text file of any barcodes used, and a primer text file. The barcodes are matched against the both ends of each sequence for assigning sequences to barcode names with no mismatches allowed to the barcode sequences. Primer sequences are matched against each sequence with no mismatches allowed. Each sequence is compared against the immediate 5’ and 3’ flanking sequences to identify the STR loci. Regular expression patterns are used to identify individual STR alleles using the known alleles from the NIST web site (http://www.cstl.nist.gov/strbase/) plus novel alleles identified by this project (Table 1). Novel alleles are named based on current nomenclature recommendations [7, 8]. A summary analysis line is included in the details report for each sequence characterized. The STR alleles and counts reports summarize the observed allele calls for each sample (identified by barcode). The Summary report includes multiple reports that characterize results by sequencing primers. This program characterizes approximately 100,000 sequences in 3 minutes on a single CPU core. The find_strs.rb program works with loci (PowerPlex 16/CODIS) with complex alleles and any STR loci with simple repeat alleles.

JalView [9] was used to visualize STR sequences.

## III. RESULTS AND DISCUSSION

The Ruby find_strs.rb program was developed to assist the scientists with the analysis of the multiplexed sequence data. Known alleles from the NIST web site (www.cstl.nist.gov/strbase) were used to characterize each sequence, which eliminates the need to assemble the sequences for each locus together. The use of regular expressions to identify known alleles eliminates the need to create a long concatemer of STR alleles as a reference sequence for allele identification as done by [5].

Twelve novel alleles, not currently included on the NIST web site, were identified in the first 10 individuals. Fordyce *et al.* [1] observed a similar level of novel alleles for five STR loci. For the D13S317 locus, 6 of 10 alleles were identified as having one extra TATC repeat unit than identified by CE results. An A to T SNP in the first base of the STR 3’ flanking sequence converts the flanking AATC sequence to match the TATC repeats. This SNP has been previously reported by [10] for the 10’ allele; this allele would be named 10(D1A->T) by the DNA Commission of the International Society of Forensic Genetics (ISFG) [11]. The 10 allele is [TATC]_10_[AATC]_2_ and the 10’ allele is [TATC]_11_[AATC]_1_. This SNP appears to be common; we have detected unreported alleles for 11’, 12’, 13’, and 14’ (Table 1) with multiple observations of these novel alleles. In addition to the D13S317 locus, a G to T SNP in the fourth base of the 3’ flanking sequence of D5S818 converts the flanking AGAG sequence into an AGAT repeat unit in a novel 13(14) allele. For one individual, both the 13 and 13(14)(novel) D5S818 alleles were identified by sequencing, which converts that individual from homozygous to heterozygous. The novel D5S818 13(14) allele is also coupled with a novel T to C SNP (U13T->C) in the 5’ flanking sequence (<u>T</u>GTAATATTTTGA to <u>C</u>GTAATATTTTGA) in multiple individuals. With a marker SNP on each side of the D5S818 STR allele, we can identify 29 of 254 (11.4%) D5S818 sequences for this individual as chimeric (likely arising from PCR artifacts from extension of incomplete PCR products followed by amplification). The chimeric sequences were proportional to the number of sequences for each strand. Chimeric sequences have been previously observed [12, 13] and represent a minority of the available sequences; allele calls can be determined from the majority, non-chimeric sequences. Finally, novel 14’ and 15’ alleles were discovered for the D8S1179 locus (Table 1). Shifting from sizing to sequencing STR alleles will provide additional information for resolving different sequences with identical sizes, while also identifying possible SNPs in the 5’ and 3’ flanking sequences. For example, an individual with a D3S1358 15/15 CE profile was resolved to 15’/15’’(novel) profile by sequencing. A similar result for two D3S1358 alleles was recently reported [14]. Sequencing STR alleles enable the detection of variant alleles that are unresolved from other alleles of the same length by allele sizing approaches. Allele calling software can identify these new alleles plus provide backwards compatibility with sized alleles results.

The number of DNA sequence reads generated for each locus was consistently imbalanced, with some loci overrepresented (Amel, D3S1358, D7S820, D8S1179, and vWA) and other loci underrepresented (D18S51, Penta D, Penta E, and TPOX) (Figure 1). This sort of imbalance in a multiplexed reaction has been reported previously [15]. In addition, an imbalance in the number of reads on each strand was also observed (Figure 2). Imbalances in the number of reads from different loci cause two problems. First, fewer samples can be analyzed in a single run, because to generate enough sequence to adequately characterize the locus that produces the fewest reads, one would produce excess data at the loci that produce excess reads. Second, it imposes a limit on the maximum ratio of DNA from different contributors in a DNA mixture. Multiplex balancing is a complex task involving a combination of informatics and empirical experimentation [16]. Increasing the primer concentration for under represented loci increased the number of reads from some loci (D18S51, FGA, Penta D, Penta E, and TPOX) and no effect on the reads from other loci (CSF1PO, D13S317, D21S11, and D5S818). Additionally, while altering the primer concentration can affect the number of reads produced at those loci, this may be accompanied by secondary, unexpected effects at other loci. Reducing the primer concentrations for the overrepresented loci by half had little effect, but decreasing the primer concentration by 90% reduced the reads from D8S to 0, D7S by 95%, and TH01 by 90% with significant 2 to 4 fold increases in reads for CSF, D13S, D16S, FGA, and TPOX. Decreasing all primer concentrations did not improve the imbalance between the loci.

**Figure 1.**
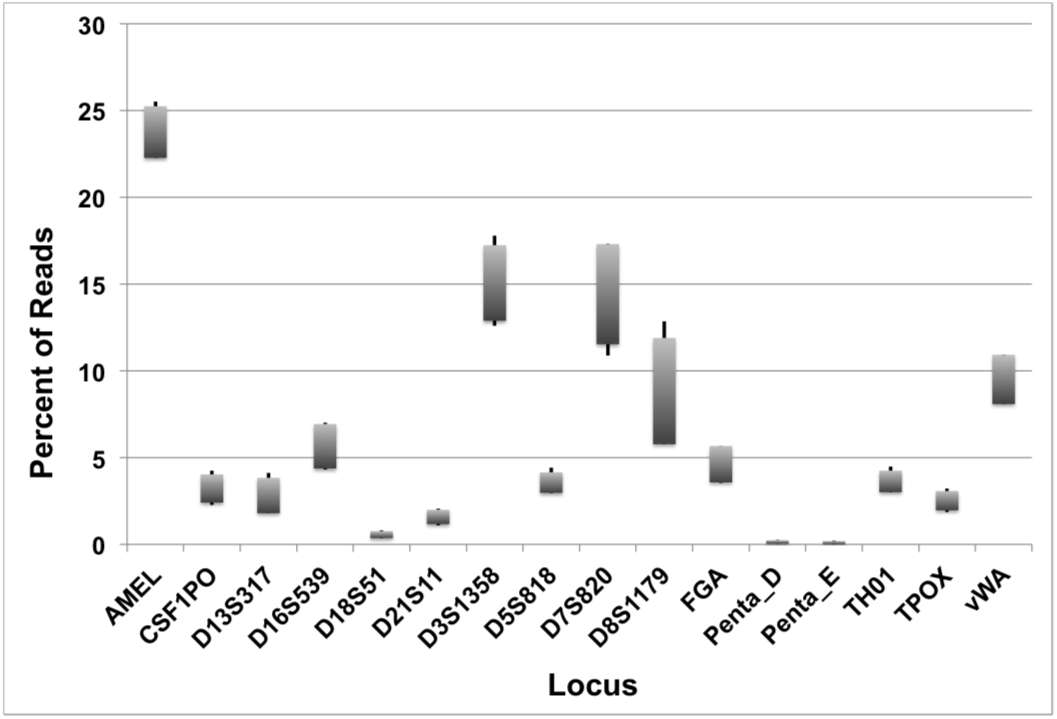
Data from 4 individuals amplified and sequenced under identical conditions showing the percent of sequencing reads attributed to each of the PowerPlex-16 loci (10 ng input DNA, 32 PCR cycles). Very little data is generated for the Penta D, Penta E, and D18S51 loci, while excess data is produced from the Amel, D3S1358, and D7S820 loci.

**Fig. 2.**
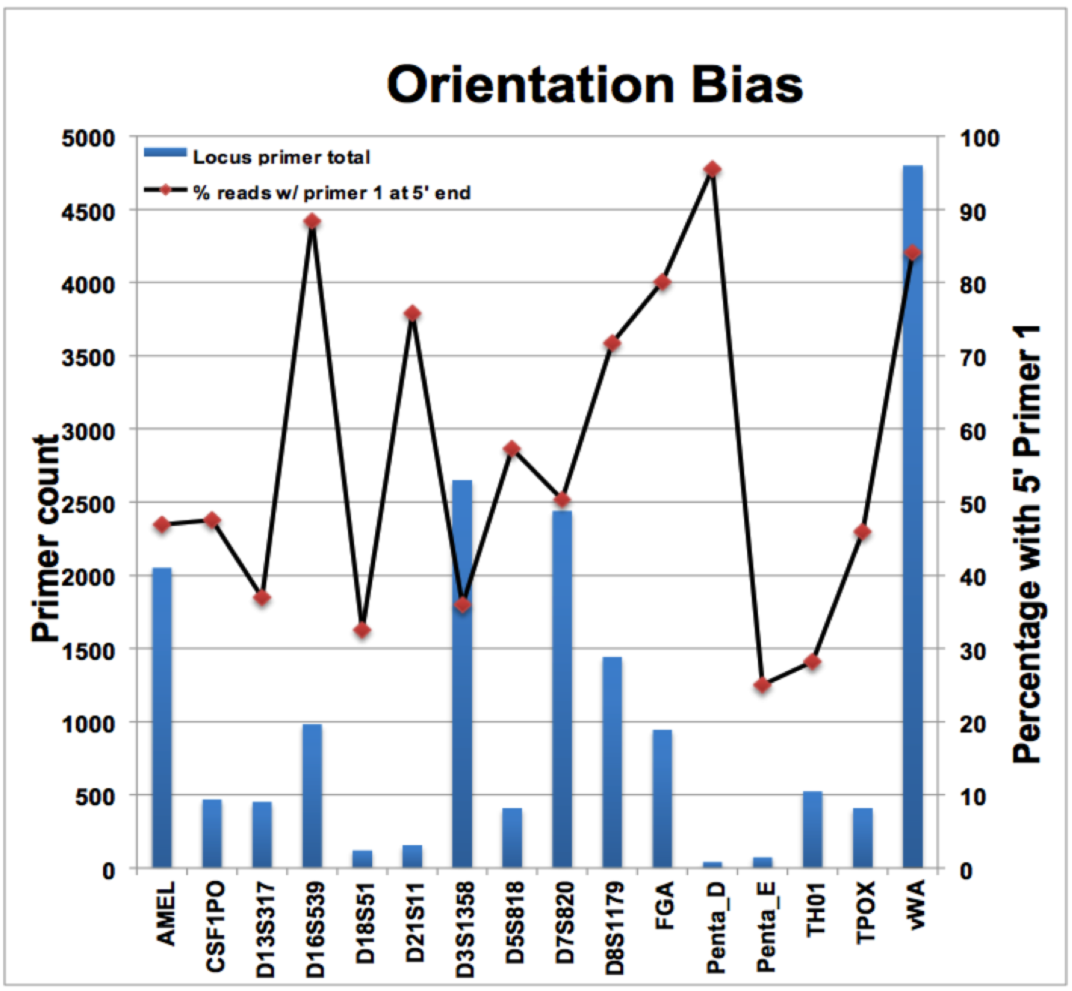
Sequencing orientation bias with counts of sequences shown in the blue bars coupled with the left side Y-axis and the percentage of these sequences shown for one of the primers by loci shown in the line graph coupled with the right side Y-axis.

The data balance between the loci can be marginally improved by reducing the number of PCR cycles from 32 6to 26 (Figure 3a), or decreasing the amount of input DNA from 10 ng to 1 ng (Figure 3b). In general, decreasing the amount of input DNA (or decreasing the number of PCR cycles) tends to decrease the number of reads from overrepresented loci (e.g. Amel, D3S1358) and increase the number of reads from under represented loci (e.g. Penta D, Penta E, D18S51).

**Figure 3.**
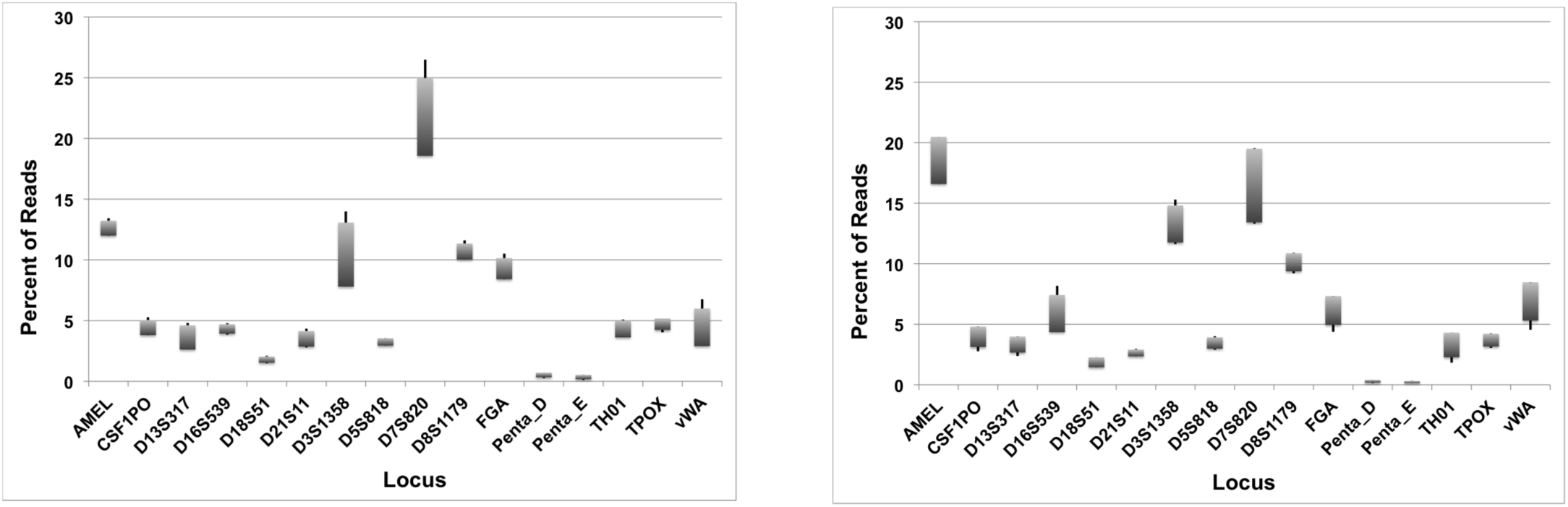
Data from 4 individuals amplified and sequenced under identical conditions showing the effect of decreasing the number of PCR cycles (10 ng input DNA, 26 PCR cycles; Figure 3a) and amount of input DNA (1 ng input DNA, 28 PCR cycles; Figure 3b) on reaction balance. In both cases, the number of reads from D18S51, Penta D, and Penta E is increased (compare to Figure 1). The number of reads from Amel and D3S1358 (high in Figure 1) decreases.

In an effort to better understand how to control the number of reads produced from individual loci, primers were generated that contained nucleotides at the 5' end that do not correspond to the endogenous sequence. Particular terminal nucleotides have been shown to affect the how well DNA is phosphorylated and/or A-tailed (See Materials and Methods section for details). Initial testing indicated that these modifications generally had the expected effect, but not in all cases. The most consistent method for decreasing the number of allele calls from a locus was to add a 5'T. Conversely, concealing 5'T residues increased the number of allele calls produced from the Penta D and Penta E loci. Combining primer alterations that increase allele calls from low abundance loci and decrease allele calls from high abundance loci led to significant improvements in reaction balance. The approach that produced the most calls from the least abundant locus was to eliminate the 5'T from the Penta D and Penta E loci, and to add 5'T residues to amelogenin, D3S1358, D7S820, and D8S1179 loci. In a perfectly balanced reaction, all loci would generate 6.25% of the allele calls. In a larger set of experiments (37 samples, 3 runs) using the modified primers and amplification conditions, the locus producing the lowest percentage of data was D18S51 which produced an average of 1.8% of the total allele calls per individual (Figure 4). While this level of balance proved sufficient for these experiments, additional optimization would be desirable for high throughput applications.

**Figure 4.**
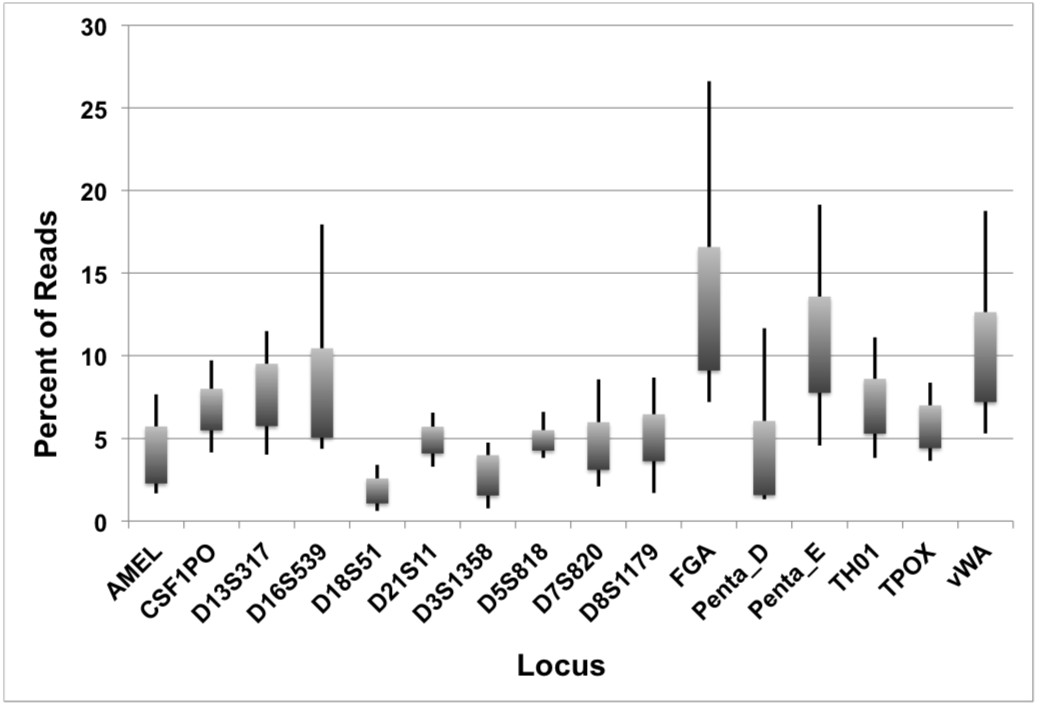
Data from 37 individuals amplified and sequenced under identical conditions (3 sequencing runs, 12 - 13 individuals per run). DNA was amplified using 5’ modified primers as described in materials and methods. These data demonstrate significant improvement in data balance (compare to Figures 1 and 3).

Ideally, HTS should produce multiple sequence reads from sequencing both DNA strands from each locus. However, for the D18S51 and FGA loci and the PowerPlex16 Penta D and Penta E loci, full-length sequences were observed for only one of the two strands for Roche 454 Jr, Ion Personal Genome Machine (PGM), and Illumina MiSeq platforms. Sequence reads for the A-rich strands, with (AGAA)_n_ and (AGAAA)_n_ repeat units, terminated in the middle of the repeats, while full-length sequences were recovered on the T-rich strands for both 454 and PGM platforms. Using the Roche GS FLX platform, Reference [17] noted that they obtained no sequences for one strand of the FGA locus. Reference [18] noted that STRs with < 10% (G+C) had consistently low coverage. The cause for these truncated or missing sequences has yet to be determined.

Amplification of repetitive regions of DNA commonly produces artifacts, referred to as stutter products. Stutter products are most often one repeat less than the true allele length (N-1), although N-2, and N+1 peaks are also commonly observed as satellite peaks in electropherograms of STR amplification products [19]. Because PCR was used to amplify STR regions prior to HTS in this study, stutter is to be expected. The level of stutter observed using PCR-sequencing is comparable to that observed using commercial, electrophoresis-based STR analysis platforms (data not shown).

Each sequencing platform has associated base calling errors. Reference [20] reports the Roche 454 GS Junior as having an average 0.38 errors per 100 bases. In general, accurate STR allele calling is reliable when an adequate number of sequences are obtained for every locus. We have a general target of 100 to 200 sequences per locus, which should accommodate sequencing errors and stutter alleles. We find that we can accurately identify STR alleles (concordant with CE allele sizing of STRs) even with multiple sequence errors in the flanking sequences (most occur towards the 3’ end of the sequences). Pyrosequencing and Ion Torrent H^+^ sequencing are well known for homopolymer-associated indel errors in base calling with error rates increasing with the length of the homopolymer. The longest homopolymer repeat occurs in the FGA locus with a stretch of six T bases. This homopolymer is frequently called as five T bases in as many as 50 to 55% of the FGA sequences, depending upon the experiment. The find_strs.rb program was modified to detect these miscalled T_5_ sequences; these alleles are labeled with ‘-’ suffix annotation. FGA alleles are identified with an additional minus suffix to indicate the number of sequences with T_5_ alleles. In one experiment, an additional T base was called in the FGA alleles consistently at the end of the fourth (CTTT) repeat for every individual in the experiment for all alleles. Obtaining sequence data from both FGA strands would provide internal confirmatory information of base call accuracy. Graphing the lengths of sequences provides a method of comparing current generation sequencing allele calls with CE allele calls. This works well for loci for which good sequence coverage is obtained, but not for low-coverage loci, including those with AAAGA repeats where only one strand is successfully sequenced.

The DNA forensics community is evaluating and likely moving towards sequencing both STRs and SNPs. HTS methods may prove useful for not only for forensic applications in human identification, but also kinship determination, mixture analysis, biogeographic ancestry analysis, and prediction of externally visible traits. However widespread acceptance of HTS for forensic applications may require that the data generated include CODIS allele call capabilities that are backwards compatible with existing DNA databases (NDIS). The current study describes methods for producing more balanced data from CODIS/PowerPlex-16 loci, and software that provides automated and accurate STR allele calling from HTS data.

